# Evolutionary potential of the monkeypox genome arising from interactions with human APOBEC3 enzymes

**DOI:** 10.1101/2023.06.27.546779

**Authors:** Brenda Delamonica, Liliana Davalos, Mani Larijani, Simon J Anthony, Jia Liu, Thomas MacCarthy

## Abstract

APOBEC3, an enzyme subfamily that plays a role in virus restriction by generating mutations at particular DNA motifs or mutational “hotspots,” can drive viral mutagenesis with host-specific preferential hotspot mutations contributing to pathogen variation. While previous analysis of viral genomes from the 2022 Mpox (formerly Monkeypox) disease outbreak has shown a high frequency of C>T mutations at T<u>C</u> motifs, suggesting recent mutations are human APOBEC3-mediated, how emerging monkeypox virus (MPXV) strains will evolve as a consequence of APOBEC3-mediated mutations remains unknown. By measuring hotspot under-representation, depletion at synonymous sites, and a combination of the two, we analyzed APOBEC3-driven evolution in human poxvirus genomes, finding varying hotspot under-representation patterns. While the native poxvirus molluscum contagiosum exhibits a signature consistent with extensive coevolution with human APOBEC3, including depletion of T<u>C</u> hotspots, variola virus shows an intermediate effect consistent with ongoing evolution at the time of eradication. MPXV, likely the result of recent zoonosis, showed many genes with more T<u>C</u> hotspots than expected by chance (over-representation) and fewer G<u>C</u> hotspots than expected (under-representation). These results suggest the MPXV genome: 1) may have evolved in a host with a particular APOBEC G<u>C</u> hotspot preference, 2) has inverted terminal repeat (ITR) regions –which may be exposed to APOBEC3 for longer during viral replication– and longer genes likely to evolve faster, and therefore 3) has a heightened potential for future human APOBEC3-meditated evolution as the virus spreads in the human population. Our predictions of MPXV mutational potential can both help guide future vaccine development and identification of putative drug targets and add urgency to the task of containing human Mpox disease transmission and uncovering the ecology of the virus in its reservoir host.

## Introduction

Monkeypox (renamed to “Mpox” by the WHO in late 2022), the disease caused by the monkeypox virus (MPXV), was first found to infect humans in 1970 in the Democratic Republic of Congo and has been mostly contained within West Africa, with local outbreaks continuing to occur since 2017-2018 (1). Nevertheless, in 2022 the global outbreak of Mpox rose to widespread concern, causing the WHO to designate it as a global health emergency in July 2022 (2). While earlier outbreaks were exported by travelers or zoonotic spillover from endemic countries in Africa, recent studies indicated the 2022 outbreak in non-endemic countries was due to regional human-human transmission within the community (3). This suggests the virus has and will continue to spread and evolve in the human population (4). It has become important to characterize the mutational potential of the emergent monkeypox virus, especially as it relates to the activity of human APOBEC3 enzymes.

The APOBEC3 enzymes, of which there are seven in humans, act as part of the innate immune system, playing a role in restriction of viruses and transposable elements. APOBEC3s cause mutations both in single-stranded DNA (ssDNA) and in RNA in the case of some family members (5), and have been found to drive restriction of certain viruses even in the absence of deaminase potential. Although poxviruses, specifically vaccinia virus, were not thought to be affected by APOBEC3s during short-term viral infection in cell culture (6), recent studies have identified a high frequency of C to T mutations in newly sequenced genomes of MPXV (7) in a short period of time. This is alarming because DNA viruses generally mutate slower than RNA viruses (8). For example, Isidro et al. (7) compared the MPXV genome MPXV-UK_P2 (See Accession number in Supplementary Table 1) from a 2017-2018 outbreak to MPXV_USA_2022_MA001 from the multinational MPXV outbreak of May 2022. There were 46 single nucleotide mutations of which 41 (89%) occurred in the context of a T<u>C</u> hotspot (7), consistent with APOBEC activity, specifically APOBEC3 (any except APOBEC3G) which have a preference for T<u>C</u> hotspots. These single nucleotide mutations most likely occurred as the virus circulated and evolved in humans following a 2017-2018 outbreak, exposing the virus to human APOBEC3 (7). Most APOBEC3s are found in the cytoplasm, except for APOBEC3B which is exclusively located in the nucleus (9–12) and APOBEC3A which locates to both nucleus and cytoplasm (13,14). Thus, because it replicates in the cytoplasm, the MPXV genome could be exposed to any APOBEC3 with the possible exception of APOBEC3B. The tissue tropism of MPXV is extensive, and the virus is capable of replicating in most mammalian cells (15). One possible trajectory after human-to-human transmission is that MPXV first replicates at the site of entry, sometimes through respiratory droplets in the lungs and oropharyngeal mucosa, or through skin to skin contact, then spreads to local lymph nodes (16). According to the Genotype-Tissue Expression (GTEx) Portal, the top three highest median bulk tissue expression sites for APOBEC3A are whole blood, spleen, and lungs. As the spleen is a major secondary lymphoid organ, it is possible that other lymphoid organs have similar expression levels. Of the remaining APOBEC3s, APOBEC3D, F and H spleen expression is also in the top three tissues, whereas expression of APOBEC3C is much more diffuse across many cell types with the exception of brain tissue. Furthermore, APOBEC expression levels are likely to be increased further as a result of cytokine stimulation during infection (17,18). Given the variety of APOBEC expression sites, and the overlap with the tissue tropism of MPXV infection, there are many opportunities for interaction. APOBEC mutations may be deleterious to the targeted virus, but may also cause diversification and thus accelerate viral evolution, as has been suggested for both HIV (19) and SARS-CoV2 (20,21).

Previous studies by ourselves (22) and others (23), have shown that viruses evolving with possible APOBEC targeting often have a statistically significant reduction of hotspot motifs in their genomes. In our previous analyses of two human gamma-herpesviruses (Epstein-Barr virus, EBV, and Kaposi-sarcoma virus, KSHV) and an alpha-herpesvirus (herpes simplex virus, HSV1), we used the Cytidine Deaminase Under-representation Reporter (CDUR) package (24) to evaluate the APOBEC3 T<u>C</u> hotspot. Briefly, the package evaluates hotspot under-representation in coding sequences by comparing the number of observed hotspot motifs (e.g. T<u>C</u>) against a null distribution derived from sequences having the same amino acid sequence but with shuffled codons. Although several shuffling algorithms are available within CDUR, both here and in previous work we have used the “n3” algorithm that generates permutations of the nucleotides at the third codon position, while preserving both the amino acid sequence and the GC content, which can greatly affect estimates if not controlled for (22). The null distribution is thus assumed to represent the distribution of hotspot motifs which could have occurred and, thus, selection against hotspot motifs is expected to result in those motifs being under-represented with respect to the null model. More formally, the percentage of shuffled sequences with fewer hotpot motifs than the original sequence is calculated. If this estimated P-value falls on the left tail of the null distribution (e.g., P < 0.05), it indicates hotspot motif under-representation, while falling on the right tail indicates over-representation (e.g., P > 0.95). CDUR also quantifies the impact of hotspot mutations on amino acid changes (amino acid replacement potential) by simulating C>T mutations at hotspots and counting the number of these that are nonsynonymous. The number of nonsynonymous mutations is then divided by the number of hotspot motifs.

Using this measure, the original sequence is again compared to the null distribution derived from the shuffled sequences. Here, if the original sequence falls on the left tail of the null distribution (e.g., P < 0.05), then amino acid changes are less likely than expected (nonsynonymous change under-representation), whereas the right tail of the distribution (e.g., P > 0.95) indicates more amino acid changes than expected (nonsynonymous change over-representation). In previous work we found T<u>C</u> hotspot under-representation in many herpesvirus genes (22). Furthermore, simulated C>T mutations at T<u>C</u> hotspots in the same genes caused amino acid changes which suggests a prolonged coevolution between the APOBEC3 enzymes and these viruses (25). We concluded that these genes had evolved under selection by APOBEC3A or B, leading to a depletion of T<u>C</u> hotspots, particularly in synonymous sites.

Although APOBEC3s have already been strongly implicated in the MPXV outbreak of 2022 (3,7,26,27), the question of how emerging MPXV strains will continue to undergo further APOBEC3-mediated mutations remains open. Thus, we have evaluated the potential for future evolution of the MPXV genome mediated by APOBEC3. In particular, we measured and compared the APOBEC3 hotspot under-representation and amino acid replacement potential across various human poxvirus genomes to evaluate the evolutionary potential of MPXV genes under APOBEC3 selection. Interestingly, our analysis of MPXV showed many genes had more T<u>C</u> hotspots than expected by chance (over-representation) and fewer G<u>C</u> hotspots than expected (under-representation) suggesting some mechanism which may have targeted G<u>C</u> hotspots, such as prior adaptation to a G<u>C</u> APOBEC targeting motif in a former host species. In some viruses it has been found that certain viral proteins may block APOBECs activity, such as Vif in human immunodeficiency virus, HIV-1, which targets APOBEC3G (28), or BORF2 in EBV which binds to APOBEC3B (29), although a recent study found that the homolog of BORF2 in cytomegalovirus (CMV) did not inhibit APOBEC3B (30). Many gene gain and loss events have been observed as part of the adaptive evolution of poxviruses, especially in the genus of Orthopoxvirus, which includes vaccinia and variola viruses (31–33). Vaccinia viruses, for example, have been seen to increase fitness relatively quickly in vitro through a “gene accordion” model (34). While gene duplication and loss in MPXV genomes may be a major component of host evasion (35) and inverted repeats may be mutational hotspots in MPXV (26), we focus part of our analysis on the T<u>C</u> hotspot representation and amino acid replacement of inverted terminal repeats (ITR), since our analysis suggests these genes may have a higher probability of being exposed to APOBEC3 during viral replication. Our results suggest that the prior evolution of the virus has led to a greatly increased present potential for APOBEC3-mediated mutations, assuming that other selective forces do not counteract this potential. If Mpox continues to spread in the human population then the abundance of T<u>C</u> hotspots may be targeted and may drive evolutionary selection of mutants with fewer hotspots.

## Results

### Comparison of human poxviruses suggests that a subset of monkeypox virus genes will be under greater APOBEC3-mediated selection

We previously developed the Cytidine Deaminase Under-representation Reporter (CDUR) package(24). As mentioned above, our previous analyses of herpesviruses suggested that viral genes under APOBEC3 selection will evolve towards hotspot under-representation together with over-representation of nonsynonymous changes for C>T hotspot mutations (a consequence of hotspot depletion biased towards synonymous sites). This pattern of evolution defines a “footprint” of APOBEC3 selection.

As shown in Figure 1, analysis of poxvirus genomes using CDUR revealed both T<u>C</u> hotspot under-representation and amino acid replacements in four human poxvirus genomes (NCBI RefSeq versions): (1) molluscum contagiosum virus (MCV), a naturally circulating poxvirus that has unique tissue tropism for human epidermis, (2) monkeypox virus (MPXV), (3) vaccinia virus (VACV), a well-characterized poxvirus also used for the smallpox vaccine and (4) variola virus (VARV), also known as smallpox. Starting with MCV, similar to viruses such as HSV-1 that have coevolved extensively with humans, and have been under APOBEC3 selection in particular, we found that MCV (Fig 1A) has a high proportion of genes with both T<u>C</u> hotspot under-representation and high amino acid replacement. In contrast, MPXV genes (Fig 1B) show no evidence for T<u>C</u> hotspot under-representation and indeed, many MPXV genes have a significant over-representation, or lack of depletion, of T<u>C</u> hotspots (dots close to the right edge of Fig 1B). As evident from the distribution of genes along the y-axis of Fig 1B, MPXV genes also show no evidence of selection for or against amino acid replacements in hotspots. These results are consistent with MPXV not yet having evolved extensively in humans, and in particular not under selection by human APOBEC3. The results for VACV and VARV are shown in Fig 1C and 1D. Previous phylogenetic analyses have shown that VACV is closely related to MPXV (36). Although the origins of MPXV and VACV are both unclear and unlikely to be the same, the CDUR results for the two viruses are consistent with a lack of extensive coevolution with their human hosts and demonstrate a significant over-representation of T<u>C</u> hotspots for many genes. The etiology of MCV is distinct from that of MPXV or VARV. In contrast to VACV, the natural hosts of VARV are humans, so more coevolution with human APOBEC3 hotspots is expected in Fig 1D. While VARV does have significantly lower hotspot under-representation (P=0.018, t-test) and higher amino acid replacement (P=0.048, t-test) than MPXV, it does not have the same “footprint” of APOBEC3 selection as MCV. The lack of a more pronounced under-representation of T<u>C</u> hotspots in VARV may be due to the slow mutation rate of DNA viruses (8,37) or that any ongoing coevolution ended once VARV was eradicated. New strains of MPXV are clearly affected by human APOBEC3 and have high mutation rates (3) so we expect genes to evolve towards the top left corner of the CDUR plot over time, much like most MCV genes, despite the differences between MCV and VARV.

**Figure 1:**
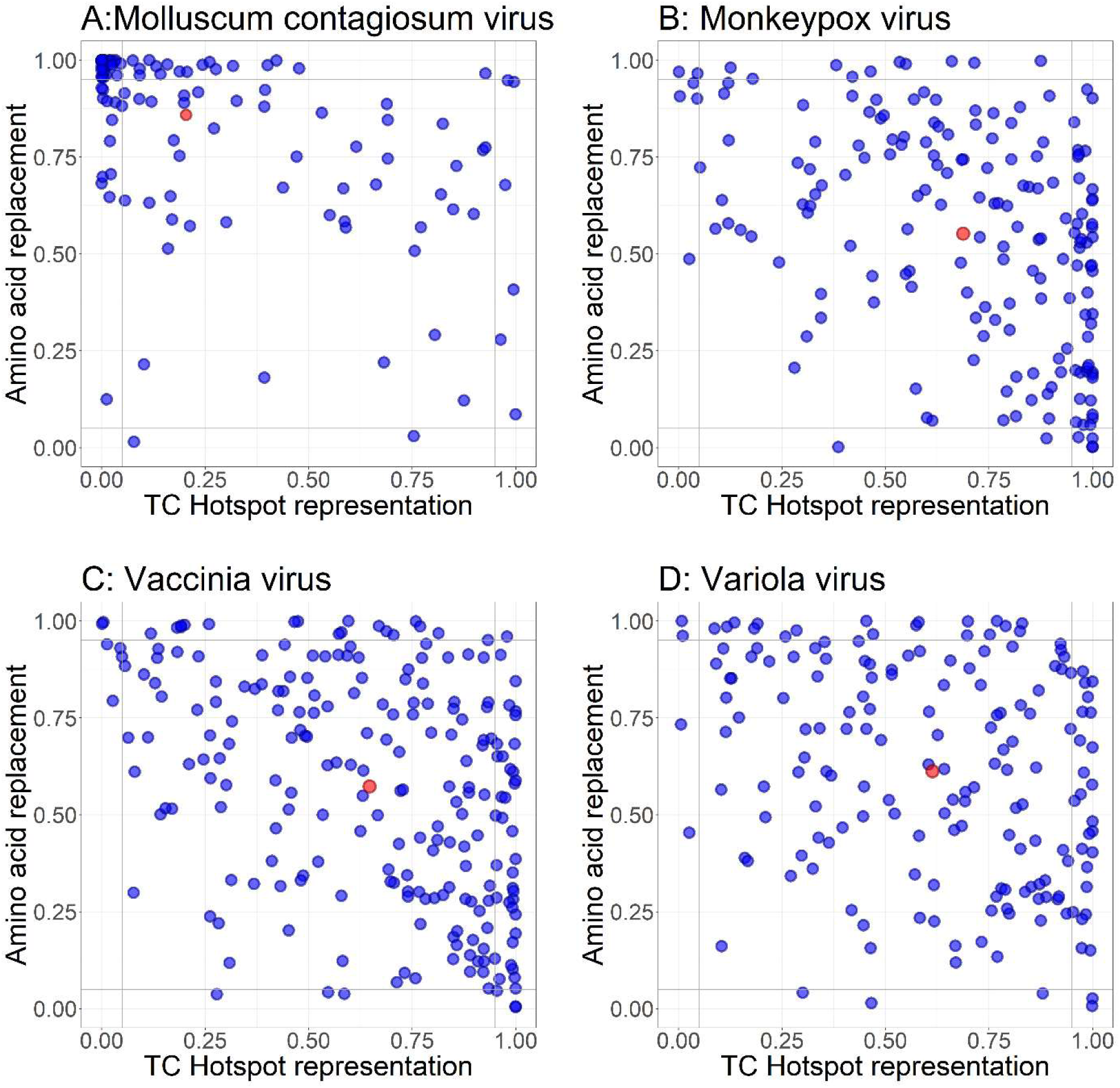
Plots of CDUR statistics (CDUR Plots) for TC hotspot under representation (horizontal axis) and amino acid replacement (vertical axis) for genes of human poxviruses. (A) Molluscum contagiosum virus (MCV), (B) Monkeypox virus (MPXV), (C) Vaccinia virus (VACV) and (D) Variola virus (VARV). Red dot indicates the mean.

To test our hypothesis that T<u>C</u> motifs in viral genomes may be depleted over evolutionary time in human hosts, we performed our CDUR analysis on a well annotated VARV strain from the 17th century (VD21, Accession number KY358055.1) and compared it to the NCBI reference sequence VARV which was collected in 1967 (38–40). Although the overall difference between the two genomes per gene is modest (Supplementary Table 9 for full list), the average shifts between genes tend slightly towards the top left corner, as expected (Supplementary Figure 1). Thus, the T<u>C</u> hotspot representation difference (VD21 minus VARV RefSeq) is positive and significant (P=0.039, paired t-test), suggesting that T<u>C</u> hotspots tend to deplete over time. The amino acid replacement difference is negative, meaning this value increases, again as expected, although the change is not significant (P=0.862, paired t-test).

### Many monkeypox virus genes show over-representation of T<u>C</u> hotspots, especially in repeat regions

Under the assumption that MPXV will continue the early trend of selection of synonymous APOBEC3 T<u>C</u> hotspot mutations (TC > TT), eventually leading to a profile similar to MCV (Fig 1A), we combined both hotspot under-representation and amino acid replacement measures to define a simple measure of mutation potential as the Euclidean distance of each gene to the top left corner of Fig 1B (i.e., hotspot under-representation=0, amino-acid replacement=1). We calculated this measure for MPXV genes using the relatively well annotated RefSeq genome. Table 1 shows the top 15 MPXV genes from Fig 1B ordered by distance to the top left corner, with genes at the top of the table having the highest distance, i.e., the highest hotspot representation and lowest amino acid replacement score (for a full list of genes see Supplementary Table 2). Many genes near the top of the list participate in immune evasion. For example, at the top of the list is a secreted chemokine binding protein (OPG001) which in VACV blocks the interaction of chemokines with cellular receptors (41). Based on the list of poxvirus genes annotated as being involved in “Host Immune Modulation” in the ViralZone database (42) we found other genes within the top 15 (not counting repeated ITR genes), including: OPG029, a homolog of VACV C6, which inhibits induction of interferons IFN-α (43) and IFN-β (44), OPG002/CrmB which binds to host Tumor Necrosis Factor (TNF) (45), and OPG019 which in VACV binds to host Epidermal Growth Factor Receptor (EGFR) to stimulate pro-viral cellular proliferation around infected cells. Interestingly, these four genes are also in the repeat regions of the genome. Among the bottom 15 genes (Supplementary Table 2) we only found one gene, OPG176, an inhibitor of the TLR4 signaling pathway in the “Host Immune Modulation” category, that has acquired a C>T mutation in the recent outbreak. While there was some enrichment (i.e., relative abundance of T<u>C</u> hotspots), at the extreme top of the list, the overall ranking of the annotated “Host Immune Modulation” genes was not significantly high or low (Wilcoxon one-sample test, P=0.2935).

**Table 1:**
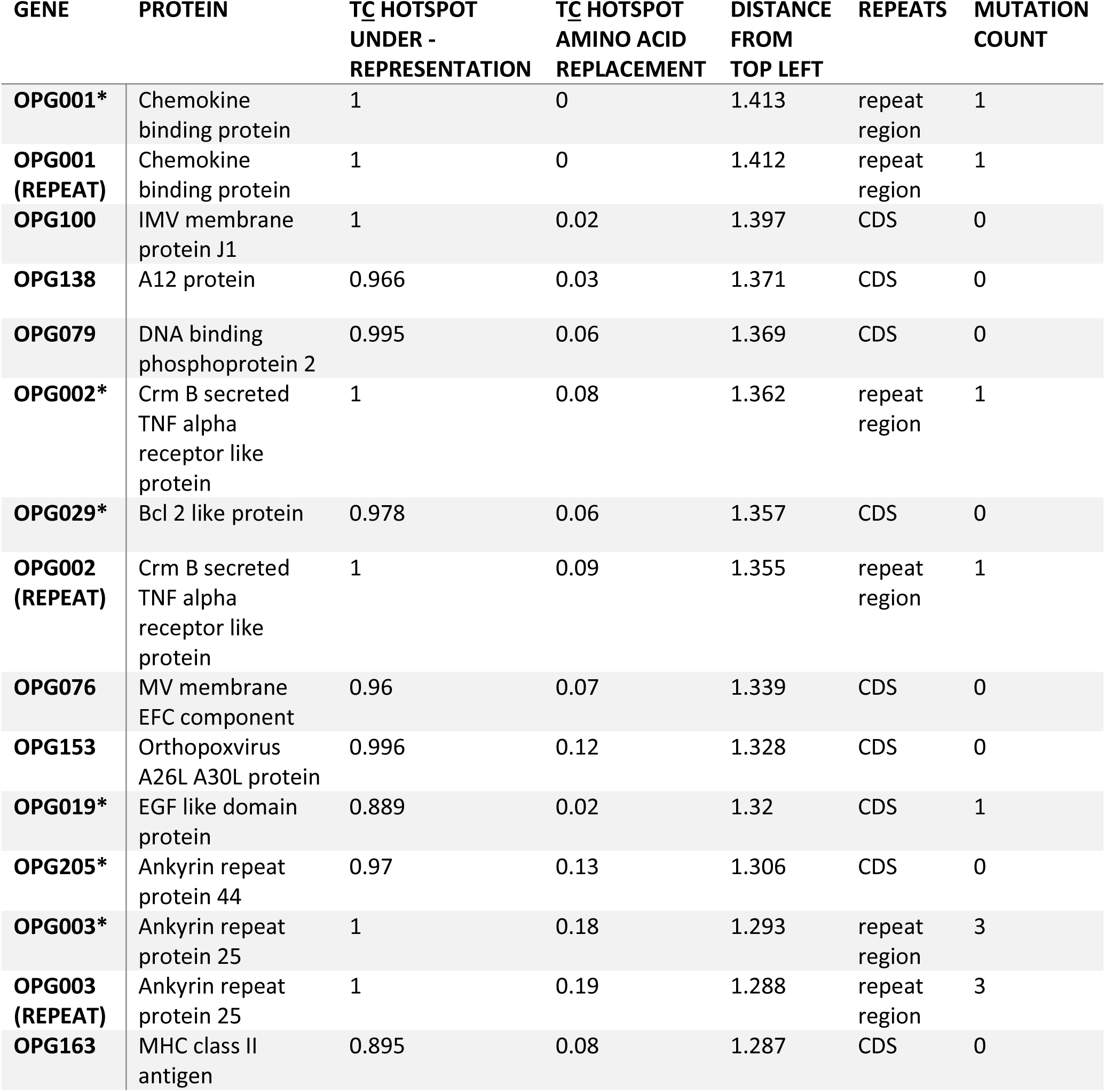
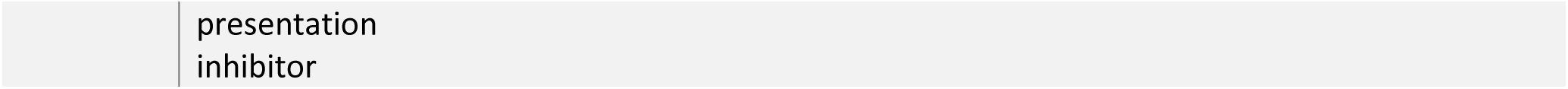
Top 15 genes in monkeypox virus genome sorted by CDUR T<u>C</u> motif result distance from the point (0,1) * = genes within 20kb of end Distance From top left = Euclidean distance of hotspot representation and amino acid replacement from point (0,1) of CDUR plot

| GENE | PROTEIN | TC HOTSPOT<br>UNDER -<br>REPRESENTATION | TC HOTSPOT<br>AMINO ACID<br>REPLACEMENT | DISTANCE<br>FROM<br>TOP LEFT | REPEATS | MUTATION<br>COUNT |
| --- | --- | --- | --- | --- | --- | --- |
| **OPG001\*** | Chemokine binding protein | 1 | 0 | 1.413 | repeat region | 1 |
| **OPG001 (REPEAT)** | Chemokine binding protein | 1 | 0 | 1.412 | repeat region | 1 |
| **OPG100** | IMV membrane protein J1 | 1 | 0.02 | 1.397 | CDS | 0 |
| **OPG138** | A12 protein | 0.966 | 0.03 | 1.371 | CDS | 0 |
| **OPG079** | DNA binding phosphoprotein 2 | 0.995 | 0.06 | 1.369 | CDS | 0 |
| **OPG002\*** | Crm B secreted TNF alpha receptor like protein | 1 | 0.08 | 1.362 | repeat region | 1 |
| **OPG029\*** | Bcl 2 like protein | 0.978 | 0.06 | 1.357 | CDS | 0 |
| **OPG002 (REPEAT)** | Crm B secreted TNF alpha receptor like protein | 1 | 0.09 | 1.355 | repeat region | 1 |
| **OPG076** | MV membrane EFC component | 0.96 | 0.07 | 1.339 | CDS | 0 |
| **OPG153** | Orthopoxvirus A26L A30L protein | 0.996 | 0.12 | 1.328 | CDS | 0 |
| **OPG019\*** | EGF like domain protein | 0.889 | 0.02 | 1.32 | CDS | 1 |
| **OPG205\*** | Ankyrin repeat protein 44 | 0.97 | 0.13 | 1.306 | CDS | 0 |
| **OPG003\*** | Ankyrin repeat protein 25 | 1 | 0.18 | 1.293 | repeat region | 3 |
| **OPG003 (REPEAT)** | Ankyrin repeat protein 25 | 1 | 0.19 | 1.288 | repeat region | 3 |
| **OPG163** | MHC class II antigen | 0.895 | 0.08 | 1.287 | CDS | 0 |

Previous analysis suggests that APOBEC3 causes mutations predominantly during viral genome replication (46,47). For example, in the case of HIV, there is a high correlation between the gradient of APOBEC3G targeting and the amount of time the DNA minus strand remains single stranded (46). More generally, APOBEC3 may target regions close to origins of replication more efficiently since the ssDNA will often be exposed for longer there, during replication initiation. A previous study in VACV identified the inverted terminal repeat regions (ITR), at the ends of the genome, as dominant origins of replication (48). We hypothesize that if ITR genes have an over-representation of T<u>C</u> hotspots they may be more susceptible to APOBEC3 mutations than other genes with high mutation potential since ITRs can potentially be exposed to APOBEC3s for longer during viral replication. We found that six of the 15 MPXV genes in Table 1 that are predicted to be most vulnerable to APOBEC3-mediated mutations, also appeared close to one of the chromosomes ends (within 20kb of either end), including within the annotated MPXV ITRs. Indeed, among the 15 genes, three out of four unique genes (six of a total of eight ITR genes) are within the top 10th percentile of all genes, with OPG001 at the top, OPG002 at the 5th percentile and OPG003 at the 8th percentile. Furthermore, these three distinct ITR genes (OPG001-3) have a hotspot representation value of close to 1 from CDUR, suggesting that genes within the MPXV ITRs are more likely to be targeted as the virus evolves in humans. Similarly, the values for amino acid replacement are all less than 0.25, suggesting they are less vulnerable to APOBEC3-mediated mutations causing amino acid changes. Of note are the ITR genes OPG001 and OPG019 which have amino acid replacement values less than 0.05 making them even less vulnerable to amino acid changes. Considering positions further away from the chromosome ends, we found only a weak negative correlation between closeness to the chromosome ends (distance in nt of each gene midpoint to the closest chromosome end) and the distance from the top left corner of the CDUR plot for T<u>C</u> hotspots (r = -0.14, P=5.9×10^-2^). These results suggest that genes closer to chromosome ends may evolve quicker in the future via APOBEC3-induced mutations, and that this effect will be strongest for genes in the ITRs. As with HIV, this effect may be a result of ITR genes at chromosome ends having a longer exposure time of ssDNA during replication than other genes within the chromosome (46). This finding is consistent with previous publications which found higher gene gain and loss activity at the ends (35) and a higher frequency of recent mutations in the MPXV inverted repeat regions (26).

### Monkeypox virus genes exhibit under-representation of G<u>C</u> hotspots

Our previous work (22,25) has shown that under-representation statistics for different hotspots will often be negatively correlated if those hotspots are mutually exclusive. For example, strong under-representation of APOBEC3A/B T<u>C</u> hotspots may coincide with an over-representation of APOBEC3G C<u>C</u> hotspots, because any remaining C sites must be either G<u>C</u>, C<u>C,</u> or A<u>C</u>. Because in the case of MPXV we observed a moderate over-representation of T<u>C</u> hotspots, we evaluated the alternative hotspots G<u>C</u>, C<u>C,</u> or A<u>C</u>, as shown in Figure 2. We found that for G<u>C</u> hotspots (Fig 2A) there is a strong effect for both CDUR measures (hotspot under-representation, amino acid replacement over-representation). Also, the under-representation measures for G<u>C</u> and T<u>C</u> hotspots are significantly negatively correlated in MPXV genes (Pearson r=-0.35, P=1.9 x 10^-6^, Supplementary Table 3). Similarly, the amino acid replacement measures for G<u>C</u> and T<u>C</u> hotspots are also negatively correlated (r=-0.2, P=8.8×10^-3^, Supplementary Table 3). Conversely, the C<u>C</u> hotspot results demonstrate a low amino acid replacement (Fig 2B), suggesting that C<u>C</u> mutations are less likely to cause an amino acid change. As expected, amino acid replacement for G<u>C</u> and C<u>C</u> hotspots are negatively correlated (r = -0.14, P=6.1 x 10^2^, Supplementary Table 3). In Table 2, MPXV genes are ranked from shortest to greatest distance from the top left corner of G<u>C</u> (rather than T<u>C</u>) hotspot under-representation and amino acid replacement (i.e., hotspot under-representation=0, amino-acid replacement=1, see Supplementary Table 4 for the full list). The G<u>C</u> results are roughly the opposite of those for T<u>C</u> (Table 1), where the same 3 distinct genes, OPG002, OPG003 and OPG001, are within the bottom 2%, 7% and 8% of genes respectively. G<u>C</u> hotspot under-representation for the 3 ITR genes does not exceed .002, while the amino acid replacement ranges between .996 and 1. An under-representation of G<u>C</u> hotspots may indicate prior exposure to nonhuman APOBECs with different hotspot preferences than those of human APOBECs (see Discussion). These findings collectively support several of our hypotheses. To summarize, if the previous host of MPXV had an APOBEC that favors G<u>C</u> hotspots, and, because the ends of the viral genomes are exposed with higher chance during viral replication, these areas may have been targeted for G<u>C</u> depletion more than other locations of the genome. Given the negative correlation between G<u>C</u> and T<u>C</u> hotspots, this could have resulted in T<u>C</u> over-representation at the ITRs of the genome, allowing a subset of genes to be more susceptible to mutation in humans with APOBEC3s that target T<u>C</u> hotspots.

**Figure 2:**
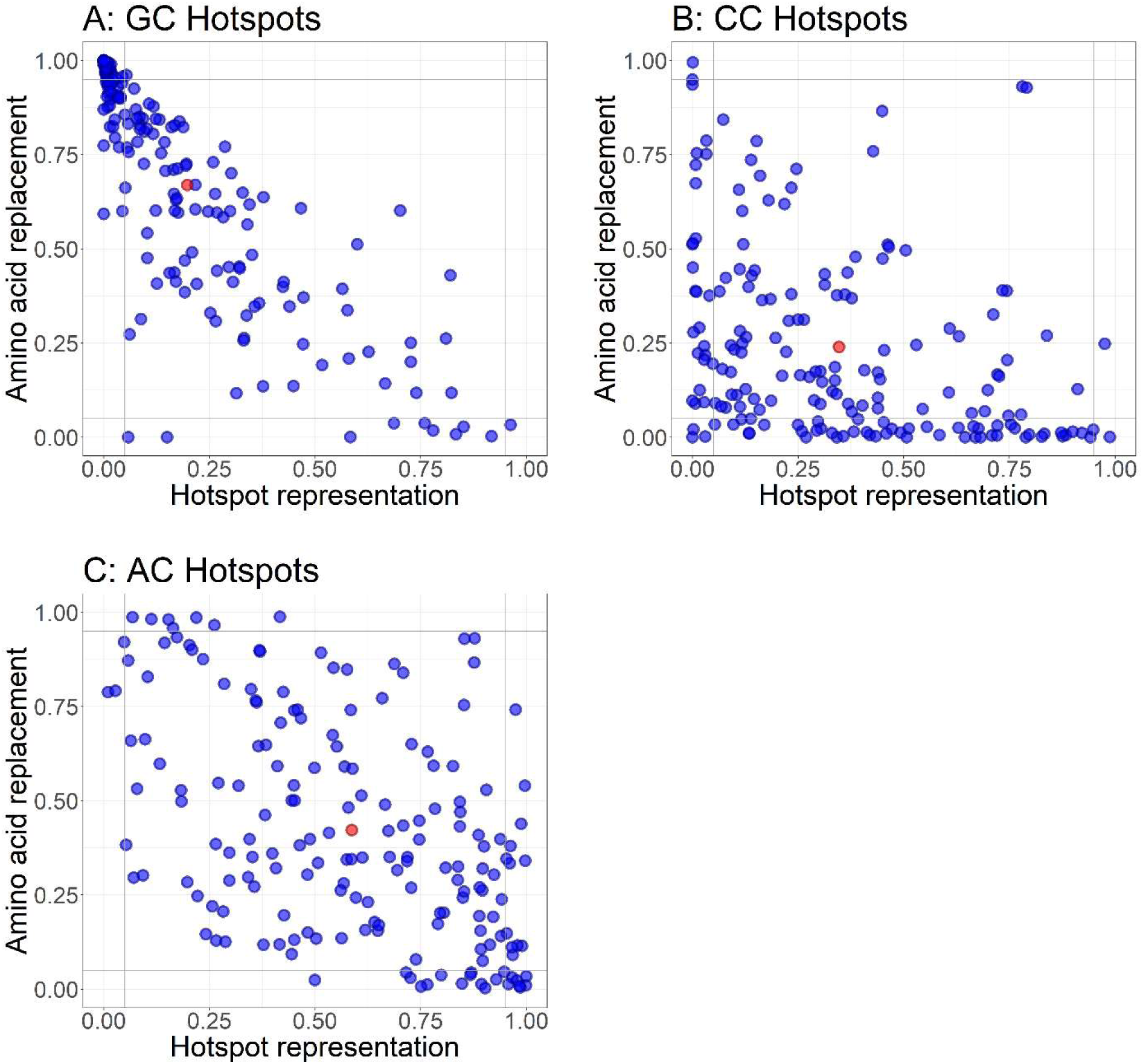
CDUR Plots for non-TC hotspots in Monkeypox virus, showing, for each gene, the hotspot under representation (horizontal axis) and amino acid replacement (vertical axis). Each plot shows results for a different hotspot: (A) GC hotspots (B) CC hotspots, and (C) AC hotspots. Red dot indicates the mean.

**Table 2:** Bottom 15 genes in monkeypox virus genome sorted by CDUR G<u>C</u> motif result distance from the point (0,1) Distance From top left = Euclidean distance of hotspot representation and amino acid replacement from point (0,1) of CDUR plot

| GENE | PROTEIN | GC HOTSPOT<br>UNDER -<br>REPRESENTATION | GC HOTSPOT<br>AMINO ACID<br>REPLACEMENT | DISTANCE<br>FROM<br>TOP LEFT | REPEATS | MUTATION<br>COUNT |
| --- | --- | --- | --- | --- | --- | --- |
| OPG002 | Crm B secreted TNF<br>alpha receptor like<br>protein | 0 | 1 | 0 | repeat<br>region | 1 |
| OPG080 | Ribonucleoside<br>diphosphate<br>reductase 2 | 0 | 1 | 0 | CDS | 0 |
| OPG105 | DNA dependent<br>RNA polymerase<br>subunit rpo147 | 0 | 1 | 0 | CDS | 4 |
| OPG002 | Crm B secreted TNF<br>alpha receptor like<br>protein | 0 | 1 | 0 | repeat<br>region | 1 |
| OPG098 | Nucleic acid binding<br>protein VP8 L4R | 0 | 0.999 | 0.001 | CDS | 1 |
| OPG117 | NTPase 1 | 0 | 0.999 | 0.001 | CDS | 0 |
| OPG200 | Bcl 2 like protein | 0.001 | 0.999 | 0.001 | CDS | 0 |
| OPG054 | Serine threonine<br>protein kinase | 0 | 0.998 | 0.002 | CDS | 0 |
| OPG189 | Ankyrin repeat<br>protein 25 | 0 | 0.998 | 0.002 | CDS | 0 |
| OPG198 | Ser thr kinase | 0 | 0.998 | 0.002 | CDS | 1 |
| OPG003 | Ankyrin repeat<br>protein 25 | 0 | 0.998 | 0.002 | repeat<br>region | 3 |
| OPG003 | Ankyrin repeat<br>protein 25 | 0.001 | 0.998 | 0.002 | repeat<br>region | 3 |
| OPG129 | Virion core protein<br>P4b | 0 | 0.997 | 0.003 | CDS | 0 |
| OPG001 | Chemokine binding<br>protein | 0.002 | 0.997 | 0.004 | repeat<br>region | 1 |
| OPG001 | Chemokine binding<br>protein | 0.002 | 0.996 | 0.004 | repeat<br>region | 1 |

This analysis focused on -1 nucleotide motifs since most human APOBEC3s have a dominant preference at the -1 nt position. However, there are exceptions, such as AID (hotspot WR<u>C</u>, W=A/T, R=A/G) which includes G<u>C</u> as a subset. To explore the relevance of the -2-nucleotide position, we evaluated the four NG<u>C</u> hotspots and found a somewhat similar trend towards G<u>C</u> amongst the three DG<u>C</u> hotspots (-2 position: A, T or G), whereas CG<u>C</u> did not show any apparent trend (Supplementary Figure 2). We conclude there is no strong preference for a -2 nucleotide in this case.

### Gene length and the potential for APOBEC3-mediated mutations

We also identified gene length as a factor for potential evolvability with human APOBEC3 (T<u>C</u> hotspots). Again, we used the distance metric (Supplementary Table 2, column “distance from left upper”) as a proxy for evolutionary potential and compared it to gene length (log10-transformed), as shown in Fig 3A1, finding a significant positive correlation (r=0.3, P=4.9 x 10^-5^, Supplementary Table 5). This is primarily due to the strong correlation between gene length and T<u>C</u> hotspot representation (r=0.42, P=1.1×10^-8^, Fig 3B1, Supplementary Table 5) as there is no correlation between gene length and amino acid replacement (r=0, P=9.8×10^-1^, Fig 3C1, Supplementary Table 5). Genes of length greater than 1.8kb (log10: ∼3.25) had very high T<u>C</u> hotspot over-representation with the exception of only two outliers, OPG210, a T-cell suppressor involved in host immune modulation, and OPG025, which also has a host immune modulation function. Eleven of the 21 longest genes are involved in genome replication, 5 genes are involved in host immune modulation and 4 in virion assembly or budding (49). Thus, longer genes may be associated with a higher likelihood of APOBEC targeted mutations (see Discussion).

**Figure 3:**
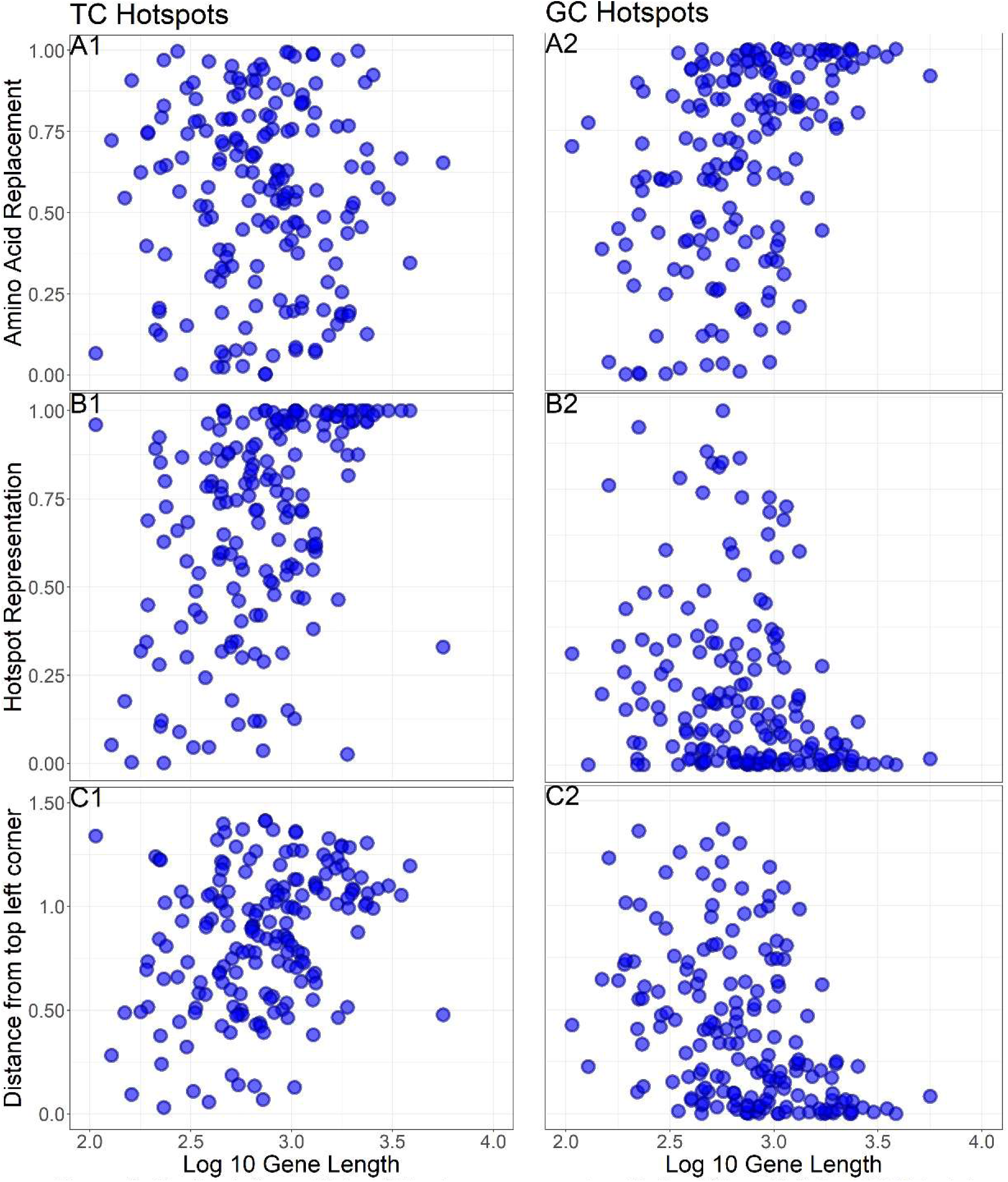
On the left are Plots of Monkeypox gene length (Iog10) vs (A1) the CDUR plot “distance from the top left corner” measure of evolutionary potential for TC hotspots, (B1) TC hotspot under representation, and (C1) amino acid replacement. One the right are equivalent plots for GC hotspots, showing Monkeypox gene length (Iog10) vs (A2) the CDUR plot “distance from the top left corner”, (B2) GC hotspot under representation, and (C2) amino acid replacement.

As mentioned above, the underlying reason for high T<u>C</u> hotspot over-representation may partially be the inverse relationship with G<u>C</u> hotspot measures (as CDUR measures should not otherwise be affected by gene length). The reason for the minimal preference for T<u>C</u> over-representation when compared to C<u>C</u> and A<u>C</u> (Fig 2) is, however, unclear. When we compared gene length with the combined measure (“distance from left upper”) hotspots, we found a significant negative correlation, as expected (r=-0.41,P=1.8 x 10^-8,^ Fig 3A2, Supplementary Table 5). We found a negative correlation between G<u>C</u> hotspot under-representation and gene length (r=-0.27, P=2.6 x 10^-4^, Fig 3B2, Supplementary Table 5), and we noticed that the 5 longest genes all had significant G<u>C</u> hotspot under-representation. Meanwhile amino-acid replacement had a positive correlation with gene length (r=0.44, P=8.2 x 10^-10^, Fig 3C2, Supplementary Table 5). Since the current MPXV strain may have originated from a previous host with APOBECs that favored G<u>C</u> hotspots, future mutations within humans may occur preferentially in larger genes, particularly those with high T<u>C</u> over-representation. As human APOBEC3s cause new C>T mutations, the number of T<u>C</u> hotspots are expected to greatly reduce, as observed in MCV. To get a better understanding of the trajectory of MPXV we considered the relationship between T<u>C</u> hotspots and gene length in MCV. Here, there is a weak negative correlation between gene length and the distance from the left upper corner of the CDUR plot (r=-0.18, P=1.9×10^-2^, Supplementary Table 6). Repeating the analysis with the two CDUR measures separately, we found T<u>C</u> hotspot representation and gene length have a very low negative correlation (r=-0.04, P=6.3×10^-1^, Supplementary Table 6) while amino acid replacement and gene length have a relatively strong positive correlation (r=0.35, P=3.7×10^-6^, Supplementary Table 6).

### Mutations in UKP2 and future evolution

A recent study (7) described mutations in the 2022 outbreak using the MPXV-UK_P2 genome (see Supplementary Table 1 for accession number) as a putative ancestor for the viruses sequenced thus far. The NCBI RefSeq MPXV and MPXV-UK_P2 were both exported from Nigeria in 2017-2018 and are closely related (7). Comparing the two genomes, MPXV-UK_P2 has 11 insertions of varying lengths (ranging from 114-2319 nt) and a total of 17 single nucleotide mutations with respect to RefSeq MPXV. Despite these differences, the two genomes are still very closely related, with an overall BLAST alignment similarity of 99.98%.

As mentioned above, previous studies reported a large proportion of TC>TT mutations ascribed to APOBEC3 (7,26,27). Isidro et al. reported a total of 46 <u>C</u>T><u>T</u>T or <u>G</u>A><u>A</u>A mutations across 27 different genes. The new mutations occurred in 3 of the 4 distinct ITR genes, as shown in Supplementary Table 7, which is again ordered by column “distance to left upper.” In particular, the genes with locus tags MPXV-UK_P2_001/190, MPXV-UK_P2_002/189 and MPXV-UK_P2_003/188 align most closely with MPXV genes OPG001, OPG002 and OPG003, and had 1, 1 and 3 mutations in each gene respectively. These 3 ITR genes are all within the 10th percentile of greatest T<u>C</u> distance from the left corner. This gives further evidence that the ITR genes are particularly susceptible to APOBEC3 mutations.

While we did not find any statistical significance between mutation count (for the new mutations) and distance from the left upper corner for T<u>C</u> hotspots, the mutation count was significantly correlated with log10 gene length (r=.70, P=4.2×10^-4^, Supplementary Table 8), as one would expect, since longer genes are more likely to be mutated. The low correlation between the CDUR measures and mutation count may be primarily due to the few mutations detected, usually one per mutated gene, and at most 4. In plotting the new mutations for MPXV-UK_P2, an association between either amino acid replacement or T<u>C</u> hotspot under-representation and new mutations (Fig 4A, compare red, mutated genes, with blue, non-mutated genes) is not immediately obvious. In contrast, the moving average of mutations over a position window of 1k nt across the genome does show two of the highest peaks at the ends, where the ITRs are located (Fig 4B). While mutation frequency (see Methods) was only marginally associated with T<u>C</u> hotspot under-representation (r=0.15, P=0.0451, Supplementary Table 8), amino acid replacement was not (r=-0.11, P=0.129, Supplementary Table 8). When we compared the combined measure (“distance from left upper”) between genes that did have a mutation against those genes that did not, we did find a significant difference (Welch two sample t-test, P=0.046). We expect this difference to become stronger as more mutations are reported including for the individual CDUR measures, amino acid replacement and hotspot under-representation. In summary, the trends observed for new (2022 outbreak) mutations suggest that T<u>C</u> hotspot under-representation and amino acid replacement will be important factors for predicting ongoing mutations of MPXV as the virus continues to circulate among humans, and that this effect is especially strong for ITR genes.

**Figure 4:**
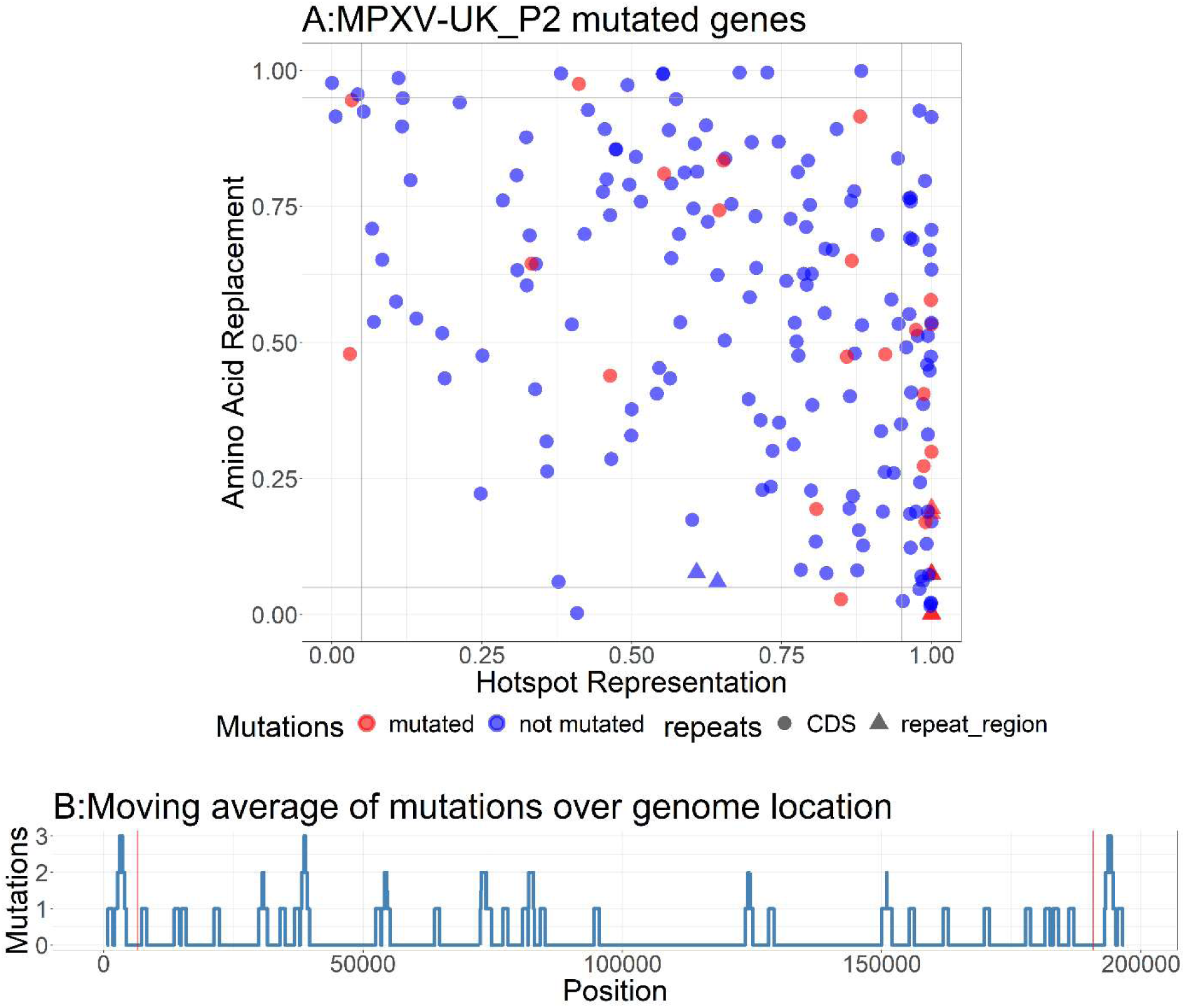
A) CDUR plot of TC hotspot under representation (horizontal axis) and amino acid replacement (vertical axis) for MPXV-UK_P2. Triangles are genes within the repeat regions (ITR) and red points are mutated genes. B) Rolling average of mutations from MPXV-UK_P2 over nucleotide position with a window of 1k nb. Red lines indicate end of repeat regions.

## Discussion

As expected for a case of recent zoonosis, MPXV genes do not exhibit signs of coevolution with human APOBEC3s such as APOBEC3A/B/H (T<u>C</u> hotspots) or APOBEC3G (C<u>C</u> hotspots). Our results show that MPXV genes have both under-representation and high sensitivity (to amino acid changes) for the motif G<u>C</u>. In part because GC and TC are mutually exclusive di-nucleotides, there is a negative correlation between the G<u>C</u> and T<u>C</u> hotspot measures (for both under-representation and amino acid replacement), which may explain why many MPXV genes have more T<u>C</u> hotspots than expected. These features may have arisen due to virus evolution in the previous host if that host had an APOBEC (or APOBEC-like) protein with a preference for G<u>C</u> hotspot mutations. Interestingly, we found that G<u>C</u> hotspot under-representation and amino acid replacement was particularly high for genes in the terminal repeat regions, which contain the most origins of replication and thus presumably expose more ssDNA to potential APOBEC mutations. Thus, we speculate that ITR genes may also have been among the most mutated, or selected, in the putative previous host as a consequence of greater APOBEC exposure. These genes were also predicted to have the highest potential for future mutation in human hosts in response to human APOBEC3 activity.

Regardless of hotspot targeting, heightened APOBEC activity would be expected to reduce the overall GC (G+C) content of a viral genome, particularly in the short-term following spillover. Yet, despite the apparent footprint of APOBEC3 on the evolution of molluscum contagiosum (MCV), the overall GC content of the MCV genome (Fig 5A) is relatively high, at ∼64% (mean per coding sequence – see Methods). In the longer term, other selection agents, such as transcriptional efficiency, which appears to be positively correlated with GC content (50), may play a larger role in changing GC content. The case of MCV suggests that codons and/or amino acids can evolve so as to reduce APOBEC3 hotspots independently of GC content. In contrast to MCV, VACV (Fig 5B), VARV (Fig 5C) and MPXV strains (Fig 5D-E) have a relatively low GC content of ∼34%. Although the discrepancy with MCV is not easily explained, it is known that certain viral families have large differences of GC content between viral species, Poxviridae being one of them (50). VACV and MPXV may have evolved a low GC content due to selective pressures in their previous host(s). For MPXV in particular, characterization of these previous selective processes will have to wait until the previous host(s) is identified. Unfortunately, APOBEC preferences have been characterized only for a few species, so this feature alone is unlikely to help identify the previous host (51). MPXV is assumed to have been transmitted relatively recently to humans from a previous, possibly rodent, host (1,52). The mouse APOBEC3 homolog (mA3) has been found to restrict retroviruses and inhibit their replication, while certain mouse viruses may have evolved mechanisms which counteract mA3 (53). In the case of mouse mammary tumor virus (MMTV), viral replication is inhibited by mA3, but G>A mutations in the integrated proviral genome are rare (54), suggesting that some mA3-mediated inhibition occurs via a deamination-independent mechanism, at least in some retroviruses. The biochemistry of deamination-independent mechanisms is still poorly understood, but it is likely that adaptation of virus genomes to deamination-independent mechanisms will lead to quite different outcomes in terms of di-nucleotide distribution, most obviously because the hotspots themselves are not directly depleted. Alternatively, the virus may have evolved in a previous organism to accumulate T<u>C</u> motifs in subregions of the genomes as an immune escape response. For example, Martinez et al. found that Epstein-Barr virus (EBV) had an over-representation of CCC motifs in particular subregions which, as the authors argued, may be acting as a “decoy” mechanism to attract AID towards those sites making it unlikely to mutate, therefore limiting mutations in more critical subregions of the genome (25). Similarly, Monajemi et al. observed HIV adaptations ex vivo and found an abundance of APOBEC3G/F hotspots in sequences which encode certain common HIV T cell epitopes. These APOBEC-mediated mutations favored CTL escape, diminishing HIV specific T cell response (19). As these examples illustrate, biased di-nucleotide distributions in certain MPXV genes may be a signature of genetic robustness in response to previous natural selection by host APOBECs.

**Figure 5:**
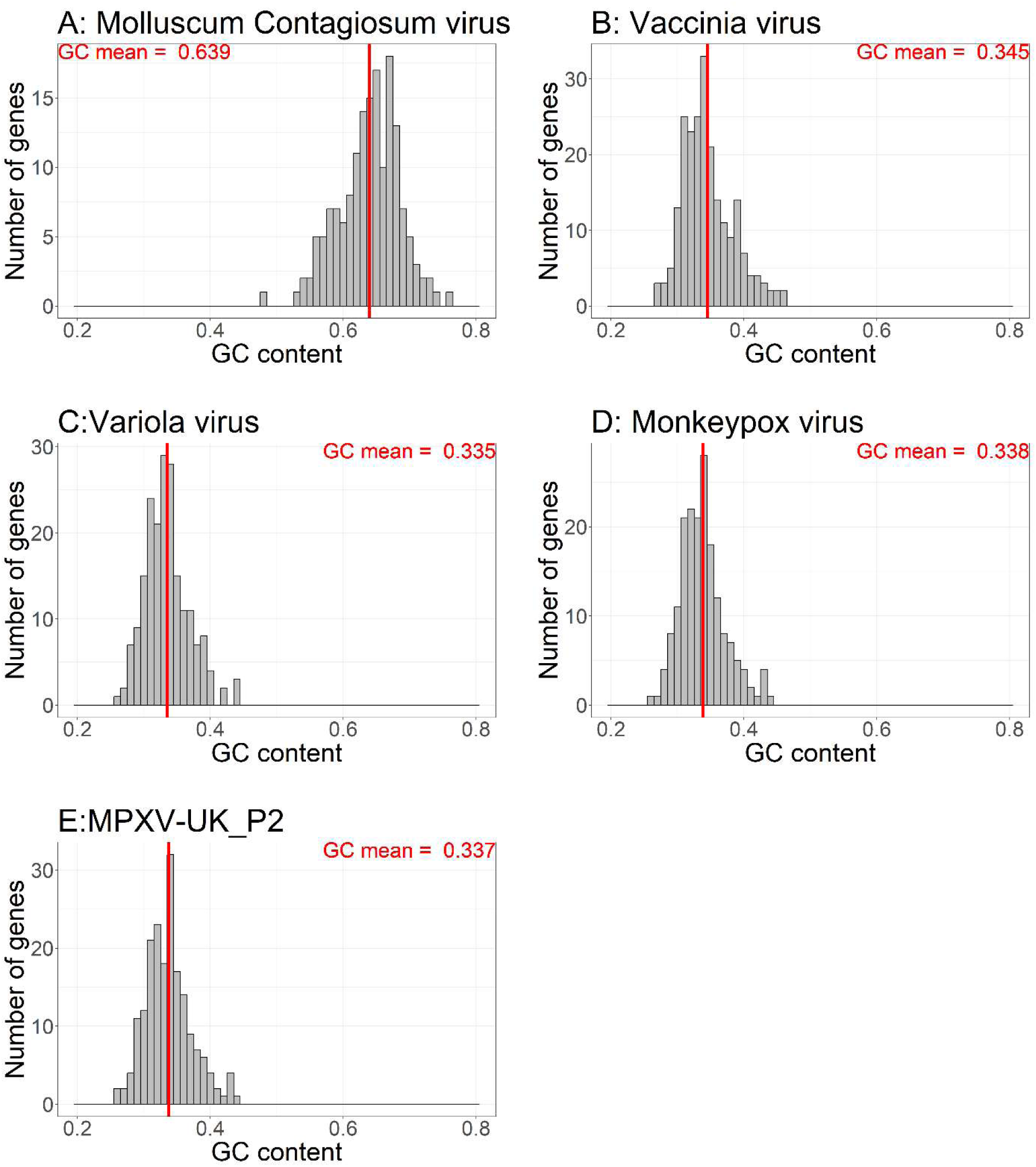
G+C content histograms and averages of (A) Molluscum contagiosum virus, (B)Vaccinia virus, (C) Variola virus, (C)Monkeypox virus, and (D)MPXV-UK_P2. Red lines indicate the average G+C content.

In primates including humans, the APOBEC3 cytidine deaminases appear to be under strong diversifying selection, which has led to the expansion and specialization (55) of the seven genes in the subfamily. Previous work has shown evidence of viral genome evolution in response to human APOBEC3s, including DNA viruses (herpesvirus, papillomavirus, polyomavirus and others), both endogenous and exogenous retroviruses and RNA viruses (coronavirus) (46). A key feature affecting the viral evolution process is the amount of time the host and viral species have had to coevolve. Well-established native species of virus in humans such as papillomaviruses, polyomaviruses and herpesviruses contain a genomic “footprint”, most obviously T<u>C</u> hotspot depletion, indicative of extensive coevolution (23,56). Our analyses reveal signatures of coevolution with human APOBEC3s (having T<u>C</u> hotspot preferences) for molluscum contagiosum virus (MCV), a native human poxvirus (Fig 1A), but not for the recently emerged monkeypox virus (MPXV). Given that poxviruses are DNA viruses that replicate in the cytoplasm, they might also be expected to be exposed to APOBEC3G, which has a C<u>C</u> (or CC<u>C</u>) hotspot preference, but we found no evidence for this in MPXV (Fig 2B) or VARV (Supplementary Figure 3), which are both very closely related with very similar hotspot distributions. However, as the virus circulates in humans, APOBEG3G may also deplete C<u>C</u> hotspots much like MCV (Fig 6). In contrast to well-established native species, viral species that have recently emerged in new host species might be expected to accumulate APOBEC3 mediated mutations as they evolve, which appears to have happened in the case of SARS-CoV-2 (20,21).

**Figure 6:**
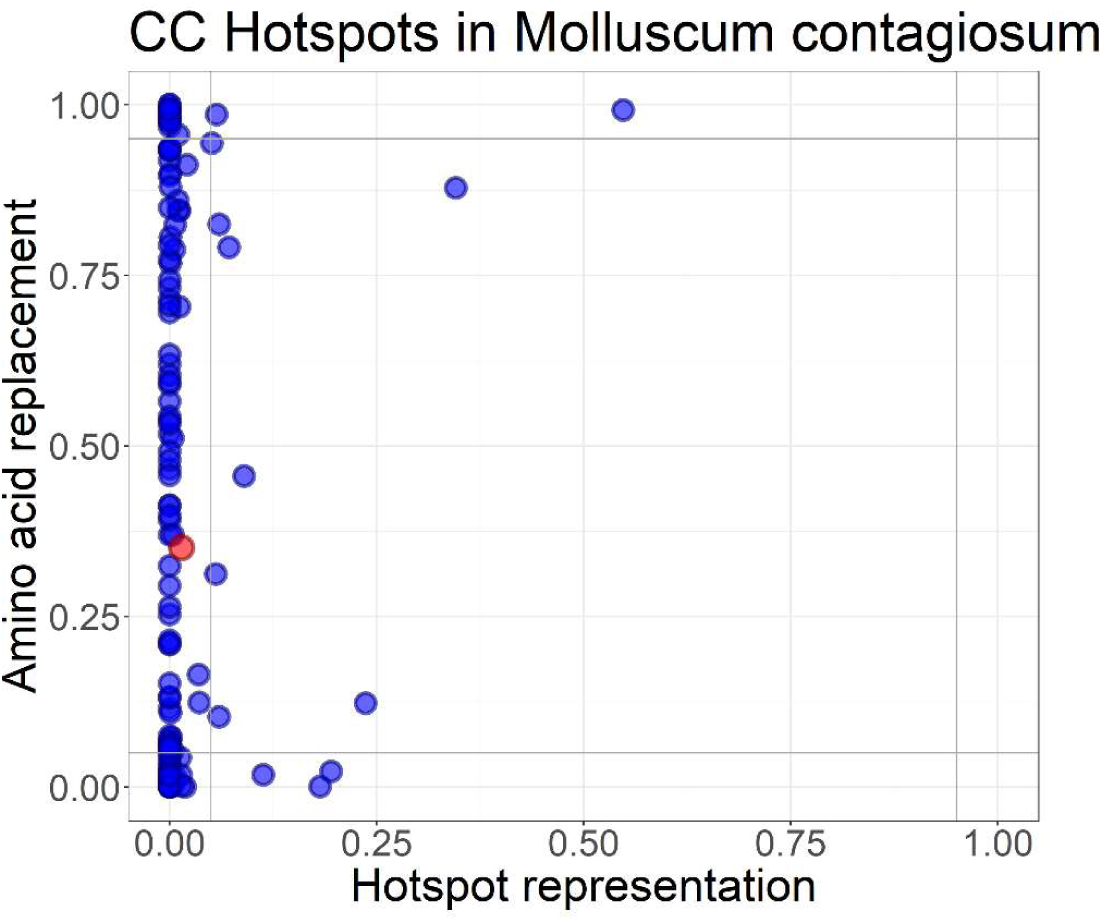
Plot of CDUR statistics for CC hotspot under representation (horizontal axis) and amino acid replacement (vertical axis) for genes of Molluscum contagiosum. Red dot indicates the mean.

We found an association between gene length and the potential for APOBEC3-mediated mutations, despite the method correcting for length by comparing with reshuffled versions of the same sequence. The simplest explanation may be that accumulation of more mutations increases the probability of one of these mutations being functionally deleterious. At the level of transcription, genes with longer transcripts may be more vulnerable to APOBEC mutations due to the presence of multiple polymerase molecules interfering with one another during transcription, which may in turn increase the exposure time of ssDNA. Recent work also suggests that ssDNA secondary structure is an important factor for deamination and can partly compensate for generating high affinity substrates even when no hotspot is present (57). Thus, although hotspot representation and amino acid replacement are dominant factors, gene length and secondary ssDNA structure may also combine in nontrivial ways to further determine APOBEC mutability. We propose to investigate in the future the impact of ssDNA secondary structure on our existing hotspot enrichment measures.

Further experimental work will be required to validate APOBEC3-mediated restriction of MPXV using cell-based assays. For example, with deep sequencing we will be able to verify whether the genes we have identified are most susceptible to APOBEC3-mediated mutations, which allow us to confirm our findings that ITRs and/or gene length are important attributes in APOBEC3 mutation potential. Other future studies could test if any of the mutations in genes we have identified change the efficacy of existing drugs or vaccines for VARV which have been used therapeutically during the 2022 Mpox outbreak (58–60). Treating Mpox with Brincidofovir remains an option. The protection of the current vaccine towards MPXV that was developed against VARV is imperfect. Tecovirimat, a pan-Orthopoxvirus inhibitor drug used for VARV, is still in clinical trials for use against MPXV and its effectiveness is not yet complete for evaluation at the time of writing (58,59,61). Tecovirimat targets the homolog of the VACV F13 gene, which produces a Palmitoylated EEV membrane protein in MPXV (62,63). Interestingly, this is one of the genes that underwent a mutation from MPXV-UK_P2 to MPXV_USA2022_MA001 (Supplementary Table 7). Gigante et al. evaluated this mutation using an infectivity assay and found that the median EC50 for attenuation of the new virus was more than twofold higher, even though the difference was not significant. Since the authors used an in vitro assay, the in vivo efficacy may turn out to be lower. While APOBEC3 may also cause other functional mutations, the correspondence between therapeutic targets and mutations can be another line of information on future treatments. Biochemical characterization of APOBECs may reveal specific targeting preferences that could be used to narrow down or identify the possible reservoir species.

In our analysis, MPXV genes predicted to mutate with highest frequency were predominantly those that have acquired most observed mutations during the current (2022) outbreak. These results suggest that MPXV, as it adapts to the human host APOBEC3s, may acquire more mutations. At the time of writing the 2022 outbreak appears to be declining, which should reduce diversification, but obviously future outbreaks that again increase diversity are possible. Also, further mutations may produce less benign strains than those currently circulating given that selection acts primarily at the level of transmissibility rather than virulence (64). While the case for containment has been made on broad evolutionary grounds before (65), our analyses outline the mutational landscape of coevolution, and have been partially validated by observed mutations.

In conclusion, we show patterns of mutation across three poxviruses with divergent host ecologies related to their exposure to human APOBEC3 to different degrees. While enrichment for T<u>C</u> hotspots in MPXV both reflects its recent emergence in humans and perhaps past adaptation in its natural host, mutations detected so far are consistent in their location and composition with mutations preferentially induced by human APOBEC3A/B/H. Although concerns surrounding MPXV appear to have abated (66) in high income countries, this pandemic continues to unfold elsewhere and its natural reservoir remains unknown. Given the great potential for further adaptation for improved transmissibility, aided in part by mutations induced by human APOBEC3, our analyses add urgency to the task of containing human MPXV transmission and uncovering the ecology of the virus in its reservoir host.

## Methods

### Data

All genome data was downloaded from NCBI using the Accession numbers in Supplementary Table 1. The tissue tropism data described in the main text were obtained from the Genotype-Tissue Expression (GTEx) Portal (https://www.gtexportal.org/home/index.html) on June 13, 2023.

### CDUR Plots

All CDUR statistics data (Figures 1, 2, 4, 6 and Supplementary Figures 1, 2, and 3; Tables 1 and 2, and Supplementary Tables 2, 4, 7 and 9) were produced by running CDUR against the coding sequences of the NCBI genomes listed in Supplementary Table 1. CDUR is available for download on https://gitlab.com/maccarthyslab/CDUR. The Supplementary data files contain the values “below_X,” hotspot representation for mutational motif X, and “repTrFrac_belowX,” amino acid replacement for hotspot X, for each gene in the given genome. Figures 1, 2, 4, and 6 plot “repTrFrac_belowX” against “below_X” using the R ggplot package.

### GC content Plots

The GC content of the four pox viruses was calculated by using the EMBOSS function geecee which computes the fraction of G+C bases found in a given sequence by finding the mean value, dividing the sum of G and C bases with the length of the entire input sequence. We ran geecee for the coding sequences of the NCBI genomes listed in Supplementary Table 1, in turn calculating the average G and C count over each genome. (https://emboss.sourceforge.net/apps/cvs/emboss/apps/geecee.html)

### Tables

From the CDUR results we calculated “distance from the top left corner” as the Euclidean distance between the point (0,1) and the location of the amino acid replacement and hotspot representation obtained from CDUR for each gene. Table 1 shows the 15 MPXV genes with CDUR results for the hotspot T<u>C</u> that are farthest away from the top left corner (0,1). Table 2 shows the 15 genes with CDUR results corresponding to G<u>C</u> that are closest to the top left corner (0,1).

In the Supplementary Tables 2, 4 and 7 we provide all measures including metadata from the NCBI genome (gene id, locus_tag, dbxref, protein id, location start and end, gbkey) as well as calculated measures for gene length and distance from chromosome ends. The repeat region was extracted from the NCBI annotations for MPXV. Gene length was calculated from the annotated NCBI data, subtracting the location end with the gene location start. Distance from chromosome end was the minimum value between the median of the gene (mid value between location start and location end) subtracted from either genome end. The columns UKP2_id in Supplementary Tables 2 and 4 and MPXV_id in Supplementary Table 7 were found by using BLAST to align the MPXV Reference genome with the MPXV-UK_P2 genome, providing a one-to-one alignment between the two genomes. The “Mutation count” column was taken from Supplementary Table 3 of Isidro et al (7) and mapped to the corresponding UKP2 genes. We calculated the mutation frequency as mutation count divided by gene length.

### Statistical Analysis

Duplicate genes in MPXV (specifically the 4 repeated ITR genes) were removed, leaving a single copy, to conduct the statistical analysis in Supplementary Tables 3 and 6, to avoid double-counting. In Supplementary Tables 3, 5, 6 and 8 Pearson correlations were calculated using the built in cor() function in R and P values were obtained using the R linear model fit function lm(). To evaluate MPXV-UK_P2 mutations we estimated P-values and correlations by removing genes without mutations. Welch two sample t-tests were obtained using the t.test() function in R. For the comparison of VD21 and VARV only paired genes were used to conduct a Welch two sample t-test, and any unmatched genes between the genomes were not included.

## Acknowledgements

Acknowledgements and Funding Sources

The study was supported by grants NIH R01AI132507 to T.M. and NFRFE-2021-00933 (Canada) to M.L., L.D. and T.M.

The study was supported by NIH R01AI139106 to J.L., a fund from the River Valley Ovarian Cancer Coalition, and a start-up by UAMS Department of Microbiology and Immunology to J.L.

