## Supplementary figures and images for "Evolutionary potential of the monkeypox genome arising from interactions with human APOBEC3 enzymes"

### Fig1.tiff

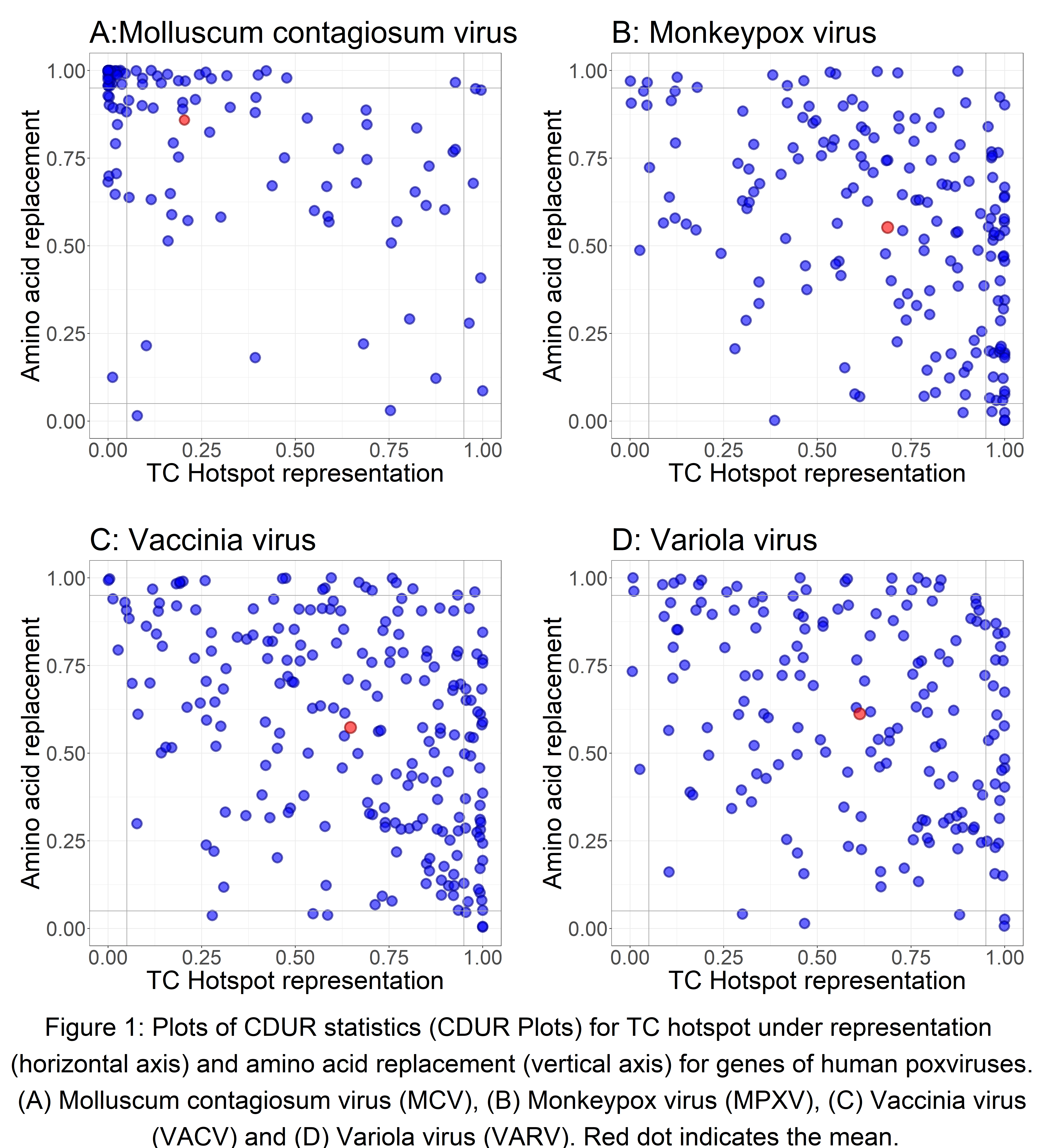

### Fig2.tiff

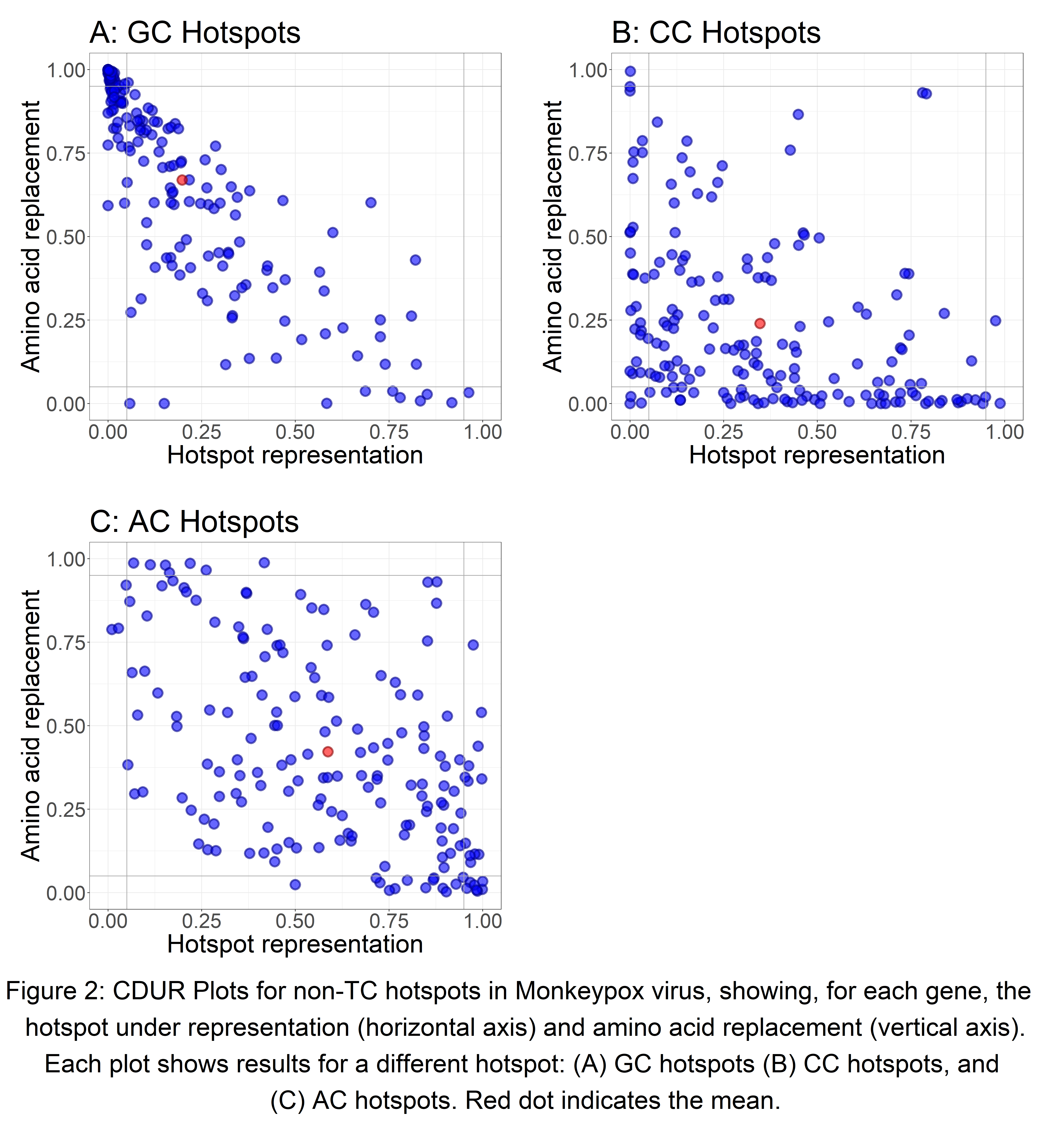

### Fig3.tiff

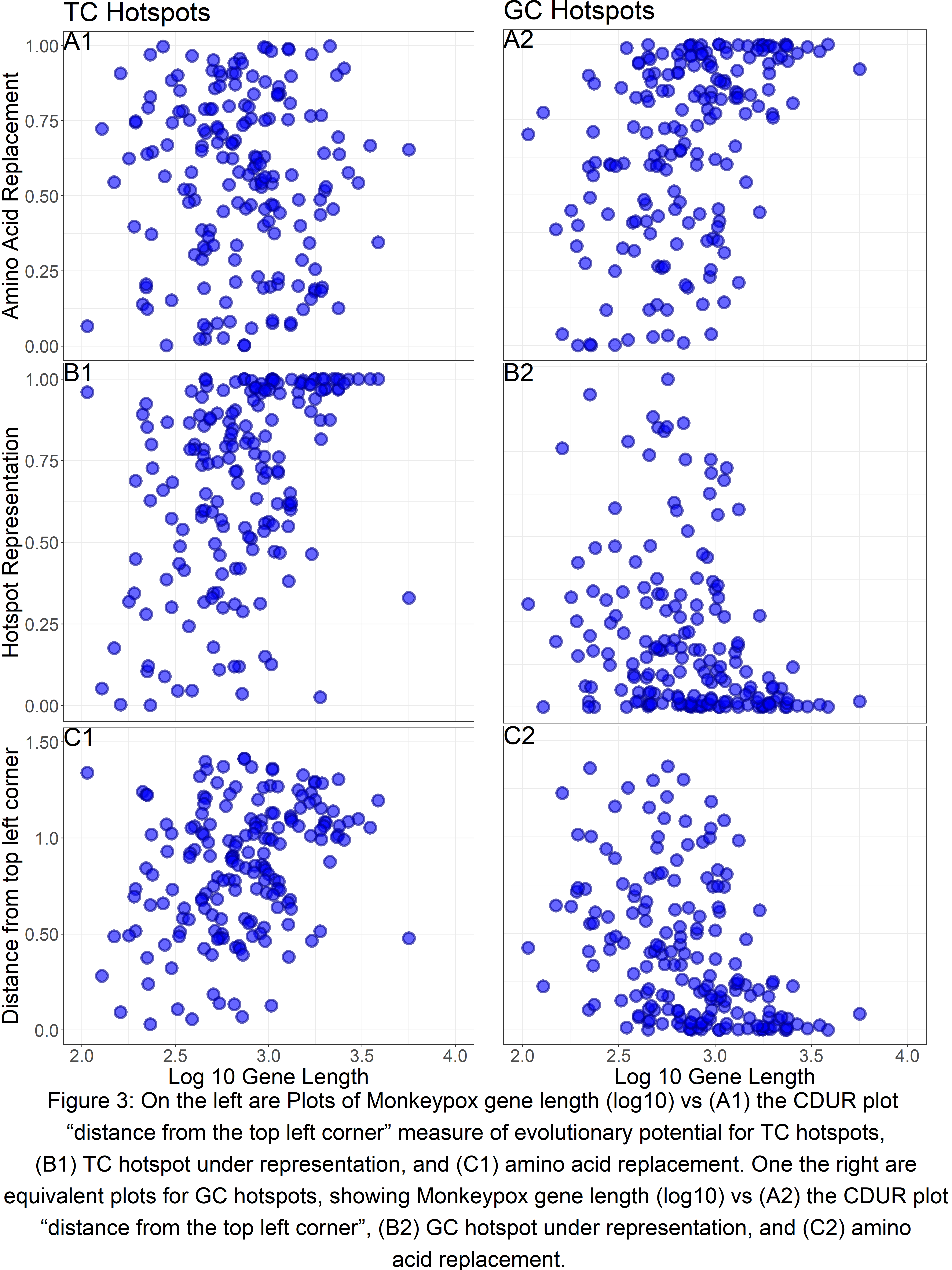

### Fig4.tiff

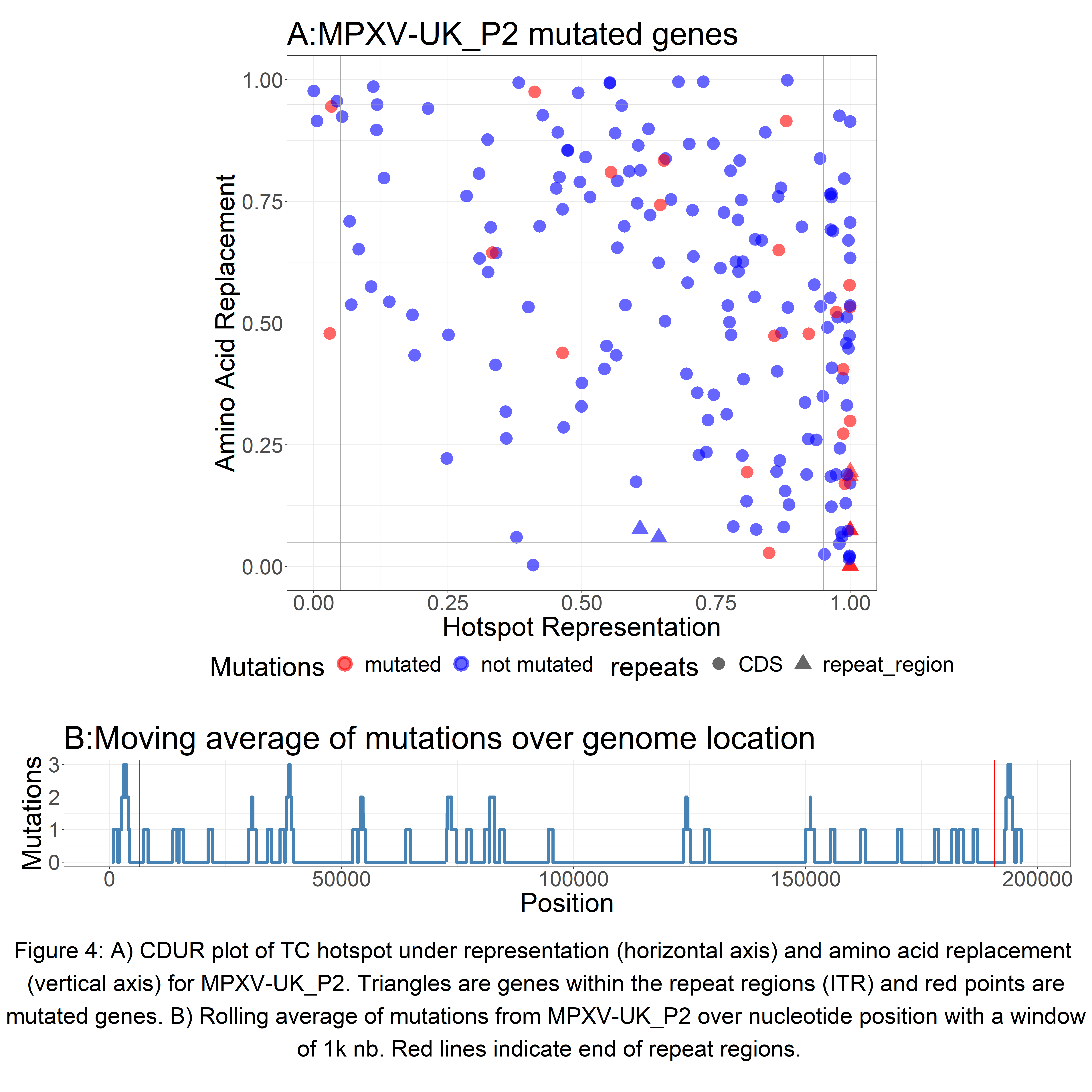

### Fig5.tiff

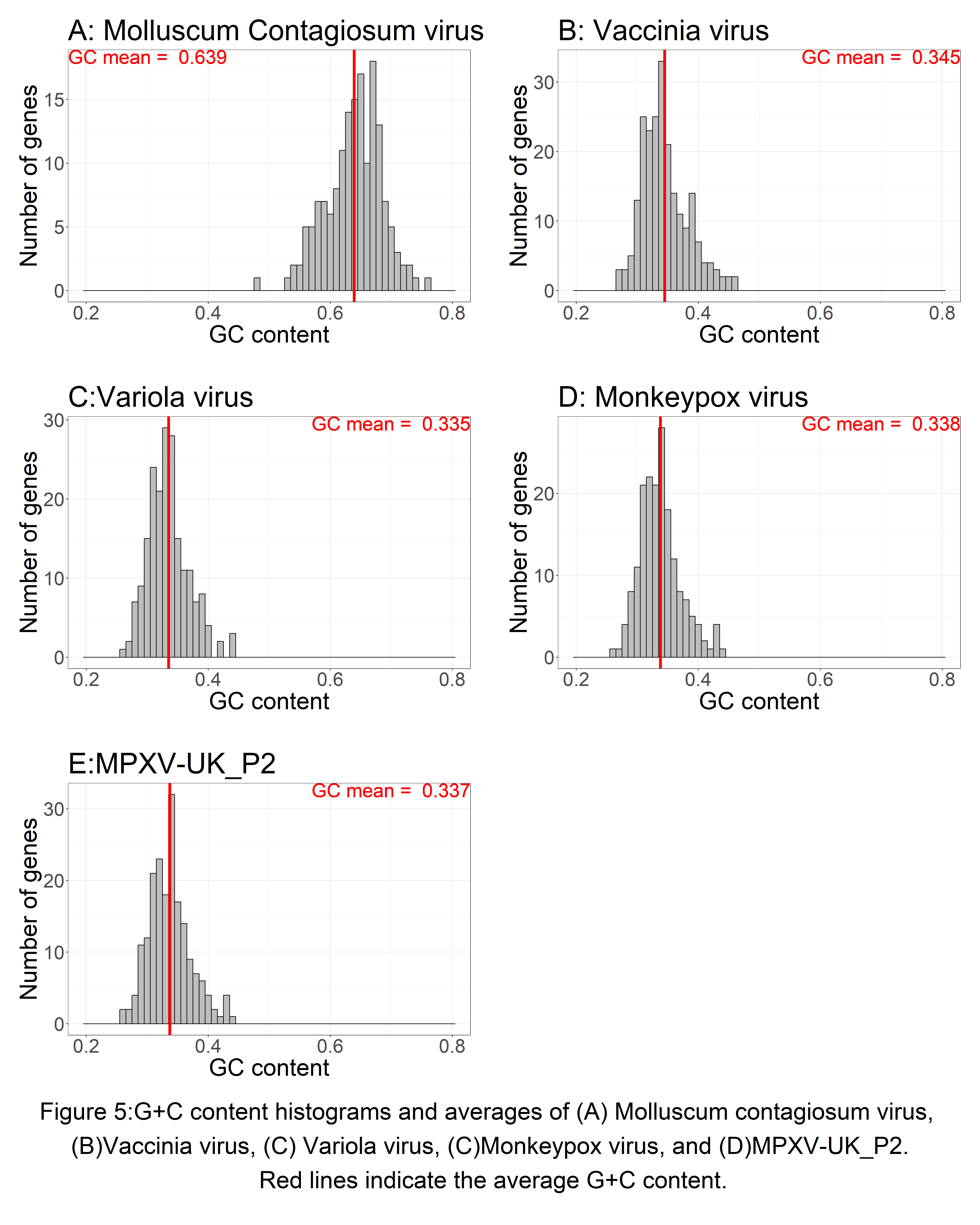

### Fig6.tiff

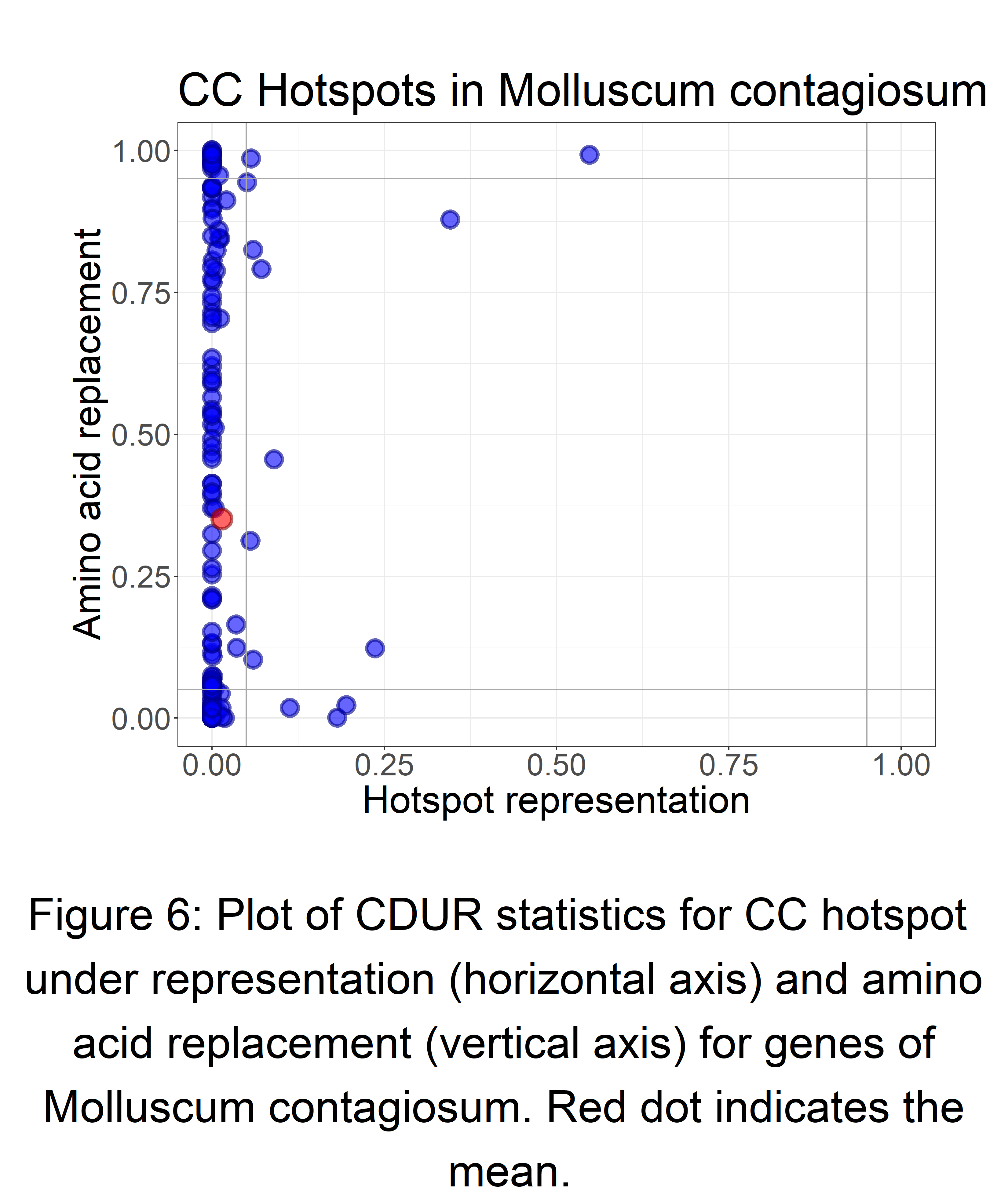

### SF1.tiff

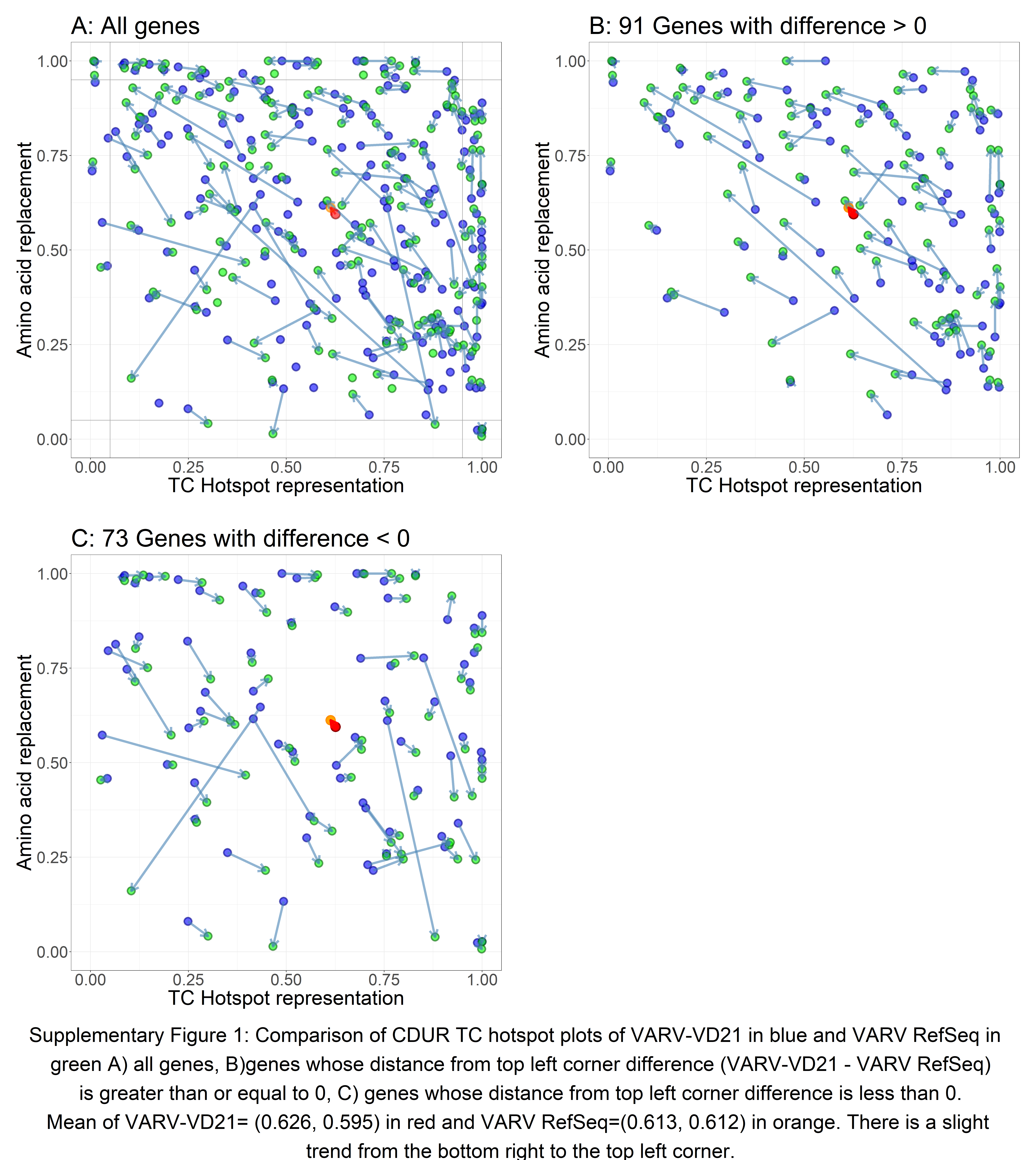

### SF2.tiff

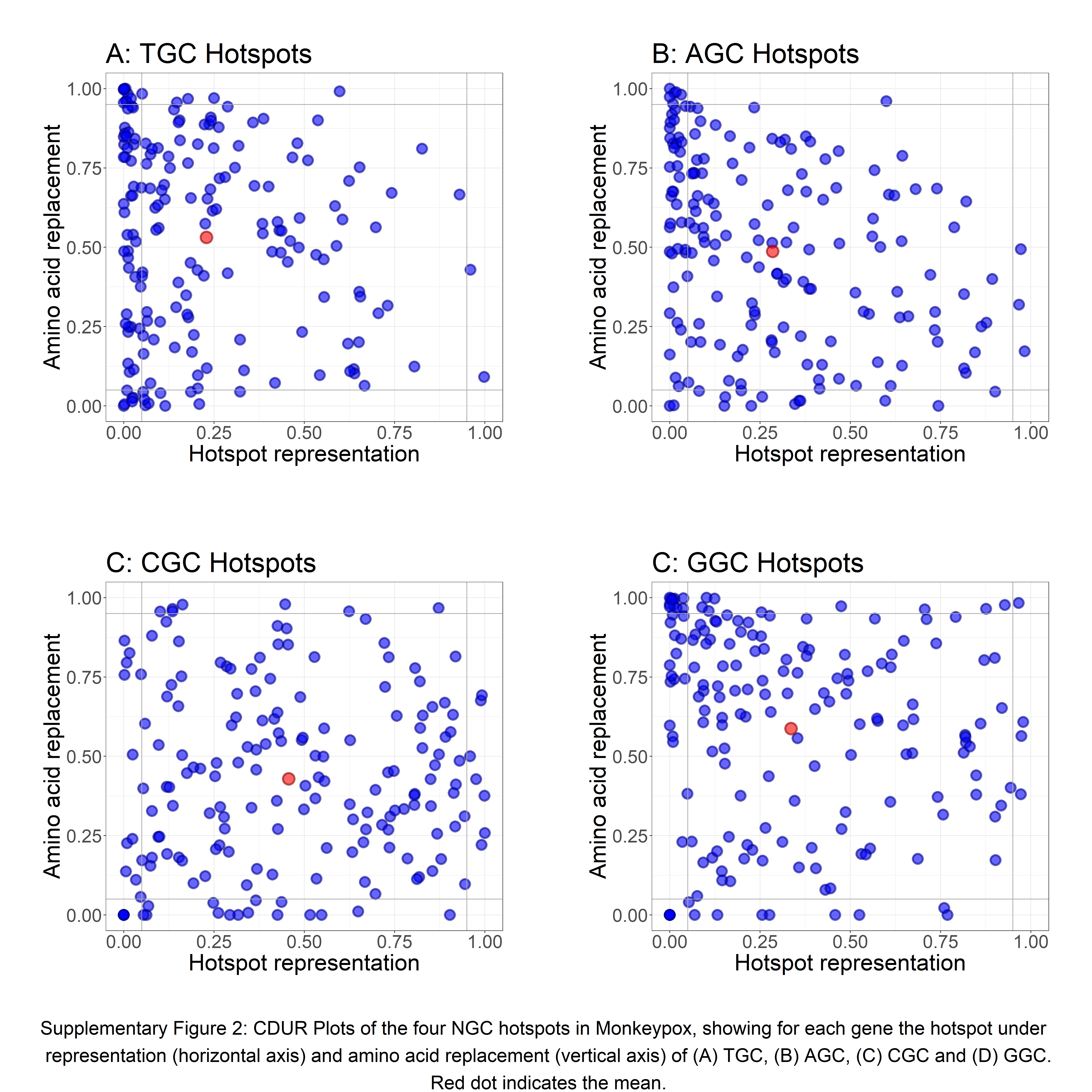

### SF3.tiff

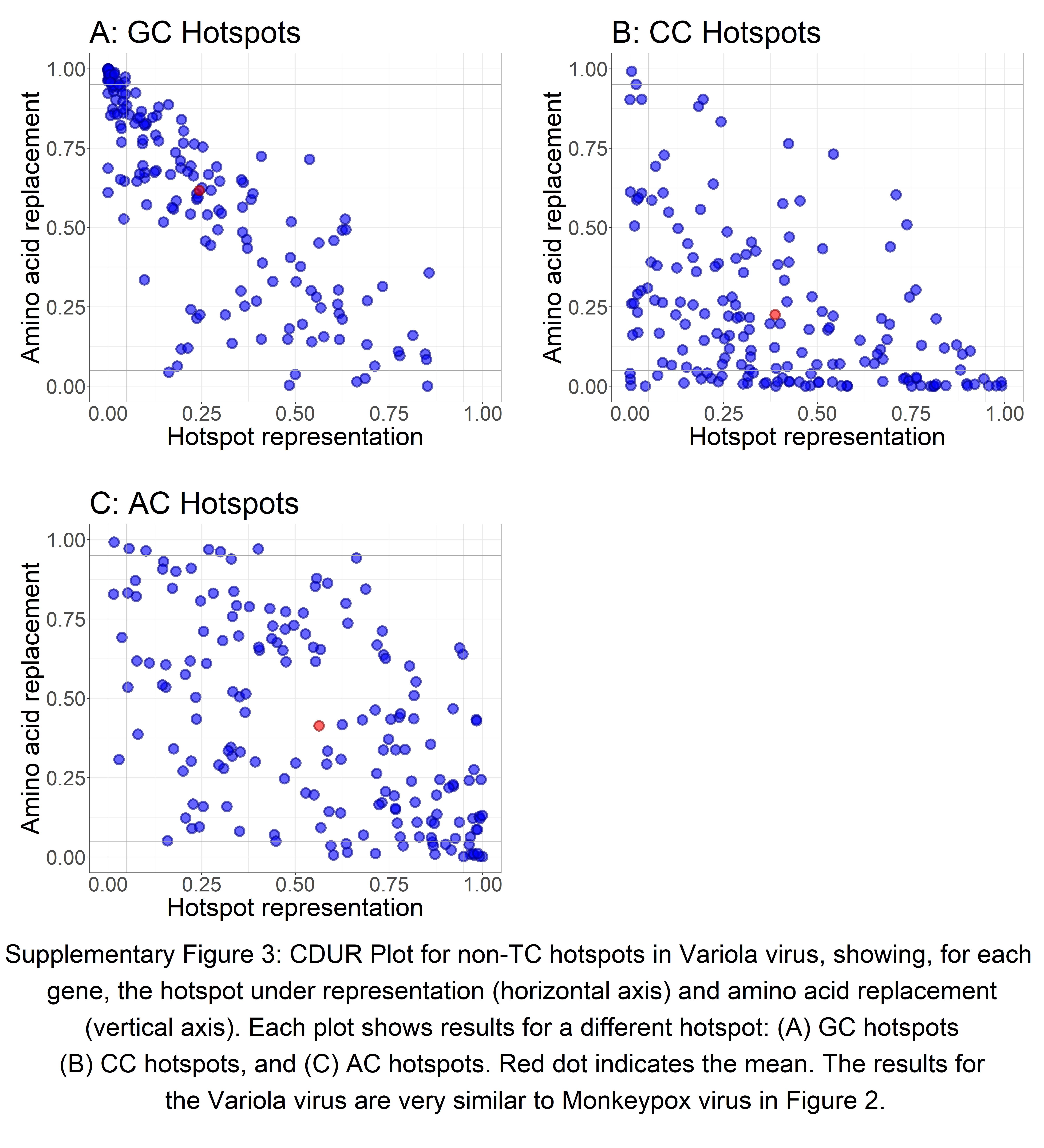
